# Metabolic mechanisms of interaction within a defined gut microbiota

**DOI:** 10.1101/250860

**Authors:** Gregory L. Medlock, Maureen A. Carey, Dennis G. McDuffie, Michael B. Mundy, Natasa Giallourou, Jonathan R. Swann, Glynis L. Kolling, Jason A. Papin

## Abstract

Metabolic interactions among species are ubiquitous in nature, and the fitness costs and benefits they impose often reinforce and stabilize them over time. These interactions are of particular importance in the human gut, where they have functions ranging from enhancing digestion to preventing (or exacerbating) infections. The diversity and sheer number of species present lead to the potential for a multitude of metabolic interactions among species to occur. However, identifying the mechanism and consequences of metabolic interactions between even two species is incredibly challenging. Here, we develop, apply, and experimentally test a framework for identifying potential metabolic mechanisms associated with interspecies interactions. We perform pairwise growth and metabolome profiling of co-cultures of strains from the altered Schaedler flora (ASF), a defined murine microbiota. We then apply our novel framework, which we call the Constant Yield Expectation (ConYE) model, to dissect emergent metabolic behaviors that occur in co-culture. Using the ConYE model, we identify and interrogate an amino acid cross-feeding interaction that is likely to confer a growth benefit to one ASF strain (*Clostridium sp. ASF356*) in co-culture with another strain (*Parabacteroides goldsteinii ASF519*). We experimentally validate that the proposed interaction leads to a growth benefit for this strain via media supplementation experiments. Our results reveal the type and extent of emergent metabolic behavior in microbial communities and demonstrate how metabolomic data can be used to identify potential metabolic interactions between organisms such as gut microbes. Our *in vitro* characterization of the ASF strains and interactions between them also enhances our ability to interpret and design experiments that utilize ASF-colonized animals. We anticipate that this work will improve the tractability of studies utilizing mice colonized with the ASF. Here, we focus on growth-modulating interactions, but the framework we develop can be applied to generate specific hypotheses about mechanisms of interspecies interaction involved in any phenotype of interest within a microbial community.

## Introduction

The structure and function of microbial communities may influence human health through a variety of means [1]. However, understanding the mechanisms governing this influence is complicated by the complexity of microbial communities. Interspecies interactions within microbial communities are essential to some benefits to human health, such as colonization resistance to pathogens [2, 3]. These interspecies interactions are often metabolic in nature, such as competition for metabolites essential for growth of pathogens [4-7]. Since metabolic interactions occur between distantly-[8] and closely-related [9] host-associated species, creating heuristics for identifying presence or absence of interactions based on phylogeny is challenging. However, knowledge of interactions among small subsets of community members has been shown to enable prediction of community assembly in larger communities, suggesting that constructing predictive models of population dynamics in complex microbial communities may be a tractable problem [10]. Thus, the ability to detect metabolic interactions may greatly improve mechanistic insight into evolution within microbial communities and associations between microbial community function and host health.

One experimental strategy used to make studies involving microbial communities more tractable is manipulation of gnotobiotic animals (i.e. animals colonized by a defined group of microbes). While this strategy greatly reduces the complexity of host-microbe studies, knowledge of the behavior of individual microbes is generally lacking, unless classical model organisms are used in place of those naturally occurring in the host-associated community (e.g. *Escherichia coli* K12, which has not inhabited a gut since it was isolated from the stool of a diphtheria patient in 1922 [11]). To improve the value of experiments performed using gnotobiotic animals, *in vitro* experiments can be performed to characterize the behavior of the species colonizing the gnotobiotic animal. The phenotyping performed via these experiments improves our understanding of these organisms, which may improve our ability to predict and interpret how they might behave *in vivo*, as in gnotobiotic animal models.

The altered Schaedler flora (ASF) is a group of 8 bacterial strains used to standardize the microbiota of laboratory mice [12]. All ASF strains were isolated from the mouse gastrointestinal tract and can be grown *in vitro*. ASF-colonized mice remain stably colonized across mouse generations and have normalized organ physiology relative to germ-free mice [12]. Although there are known differences between the immune repertoires of ASF-colonized mice and conventional mice, these differences can be exploited to test specific hypotheses (e.g. restoring the immune function with other microbes, or evaluating the role of the immune function by comparing ASF mice to conventional mice) [13, 14]. ASF mice have been used widely in infectious disease research to study *Clostridium difficile* [15], *Helicobacter bilis* [16], *Salmonella enterica* [17, 18], and *Cryptospridium parvum [19]*. Additionally, some specific pathogen-free mice (e.g. from Taconic) are initially colonized with the ASF, which has led some to theorize that presence or absence of ASF strains contributes to vendor-specific differences in susceptibility to disease [20]. Further use of gnotobiotic systems such as ASF-colonized mice could greatly accelerate discovery in microbiome research, especially if the behavior of the ASF alone is well-understood.

Previously, we performed pairwise spent media experiments using seven of the ASF strains, in which each strain was grown in a chemically-defined medium as well as the spent medium of other strains [21]. We identified cases of putative cross-feeding and competition and the effect of those interactions on growth dynamics. However, each strain was spatially and temporally separated in this study. While spent media experiments remove many of the technical and statistical complications in inferring metabolic interactions, the interactions that are possible are different in nature than those that might occur while strains are grown in co-culture.

Here, we further define the interactive potential of six of the ASF strains by performing co-culture growth experiments with all pairs of strains. We identify the influence of interspecies interactions on growth of each strain, then apply a novel statistical model for inferring metabolic mechanisms of interaction from supernatant metabolomic data. We experimentally interrogate an inferred cross-feeding interaction in which one ASF strain (*Parabacteroides goldsteineii ASF519*) produces amino acids that another (*Clostridium sp. ASF356*) consumes, confirming that the hypothesized mechanism occurs and leads to a growth benefit for the consuming strain. With this new insight, we provide a framework for mechanistic interrogation of microbe-microbe and host-microbe interactions that can be applied to any microbial community to investigate co-culture phenotypes including growth enhancement or changes in metabolite yield.

## Results

### Ecological interactions within the altered Schaedler flora

We collected in *vitro* data for growth of all pairwise combinations of 6 ASF strains (Fig 1A, n=6-9 biological replicates per strain pairing). Current taxonomic assignments [12] for these ASF strains are provided in Fig 1B. We determined the impact of co-culture on each strains’ growth by comparing monoculture abundance after 72 hours of growth to the abundance of each strain in co-culture at the same 72 hour timepoint (strain abundance determined using hydrolysis-probe-based qPCR; total density determined using OD600; at 72 hours, all strains are in stationary phase; see Materials and Methods).

**Figure 1.**
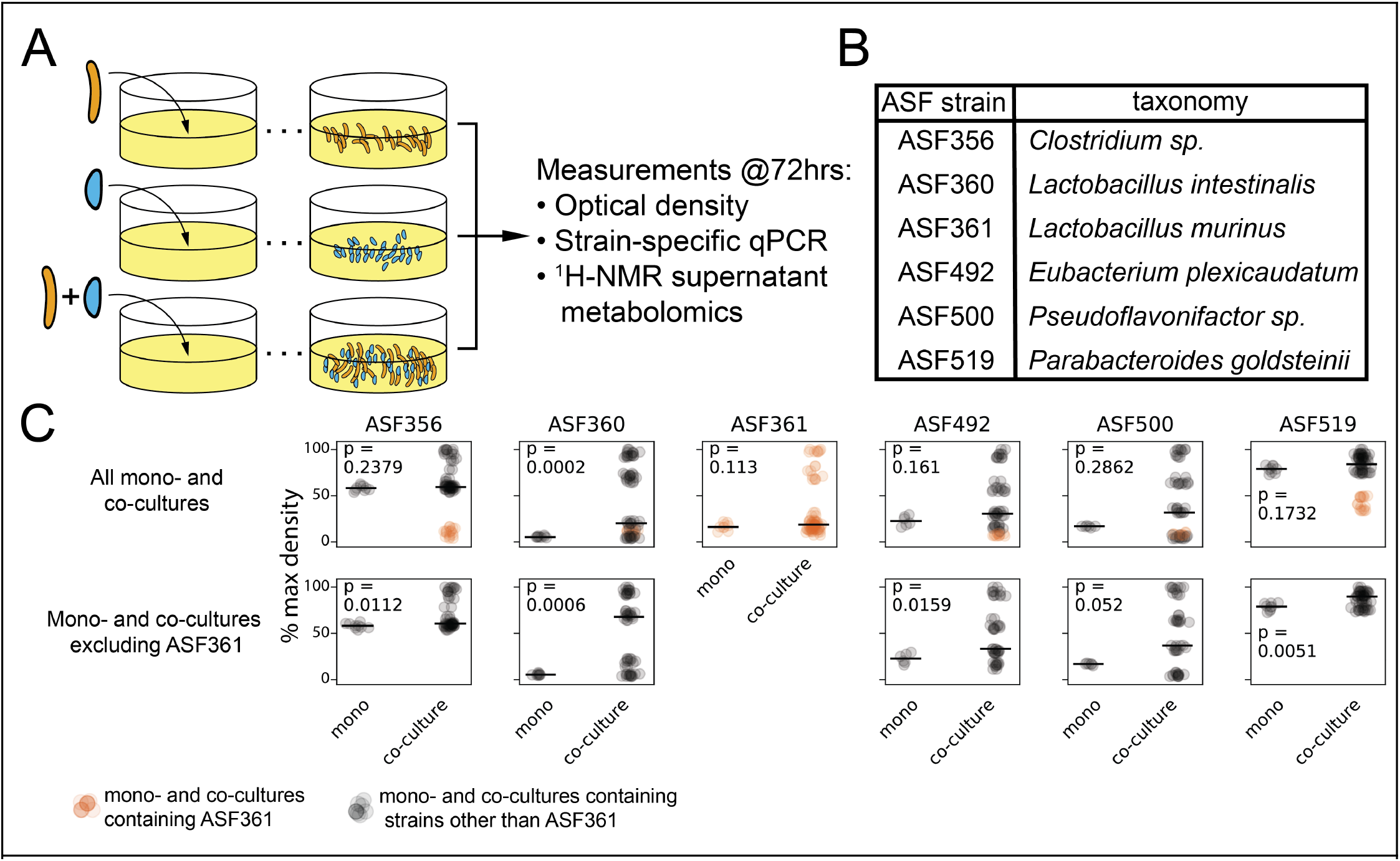
**A)** The experimental procedure for each pair of strains and measurements taken. **B)** Taxonomic assignment for the ASF strains included in this study. **C)** For each strain, the density after 72hrs of growth in monoculture (‘mono’, left in each subplot) compared to all co-cultures containing the strain in the title for the column (‘co-culture’, right in each subplot). Density determined using OD600. Max density for each subplot is the maximum OD600 from any sample within the subplot. Orange points represent samples from a mono- or co-culture group containing ASF361, while all other mono- and co-culture conditions are shown by black points. Black bar is mean of monoculture or co-culture within each subplot. Data presented in bottom row are identical to top row, but exclude samples containing ASF361. *p*-values were determined by two-sided Mann-Whitney *U* test with false discovery rate control using the Benjamini-Hochberg procedure.

For each strain except ASF360, the monoculture density was not significantly different than the density of all co-cultures containing that strain (Fig 1C, top row). However, we observed that ASF361 had a substantial negative impact on the density of all co-cultures that contained it. Removing co-cultures containing ASF361 from consideration, the co-cultures for all strains (except ASF500) grew to a higher density than the corresponding monoculture (Fig 1C, bottom row). Using DNA abundance quantified via hydrolysis probe-based quantitative polymerase chain reaction (qPCR), each strain in each pair was evaluated to determine whether a negative (-), positive (+), or neutral (0) effect on endpoint abundance occurred due to the pairing, allowing classification of the pairwise interaction with standard ecological terminology. All pairings except one, ASF356 with ASF519, had a negative impact on the abundance of at least one strain, with 0/-(amensalism), +/- (parasitism), -/- (competition), and +/0 (commensalism) being the only interactions detected (8, 4, 2, and 1 instances, respectively; data shown in Fig 2A, summarized in Fig 2B and 2C). ASF361 was present in 3/4 parasitic co-cultures and in all cases was the strain in the pairing that experienced a growth benefit. In contrast, growth of both ASF492 and ASF500 was inhibited in every condition, including in co-culture with each other. Thus, co-culture led to higher total biomass production for all co-cultures except those containing ASF361 (Figure 1C), although growth of individual strains tended to be lower in co-culture than in monoculture. These observations suggests that although these strains compete for resources, differences in resource utilization across strains, or emergent behavior in co-culture such as cross-feeding and consumption of novel substrates/metabolites, may enhance efficiency in co-culture in the absence of strong negative interactions by ASF361.

**Figure 2.**
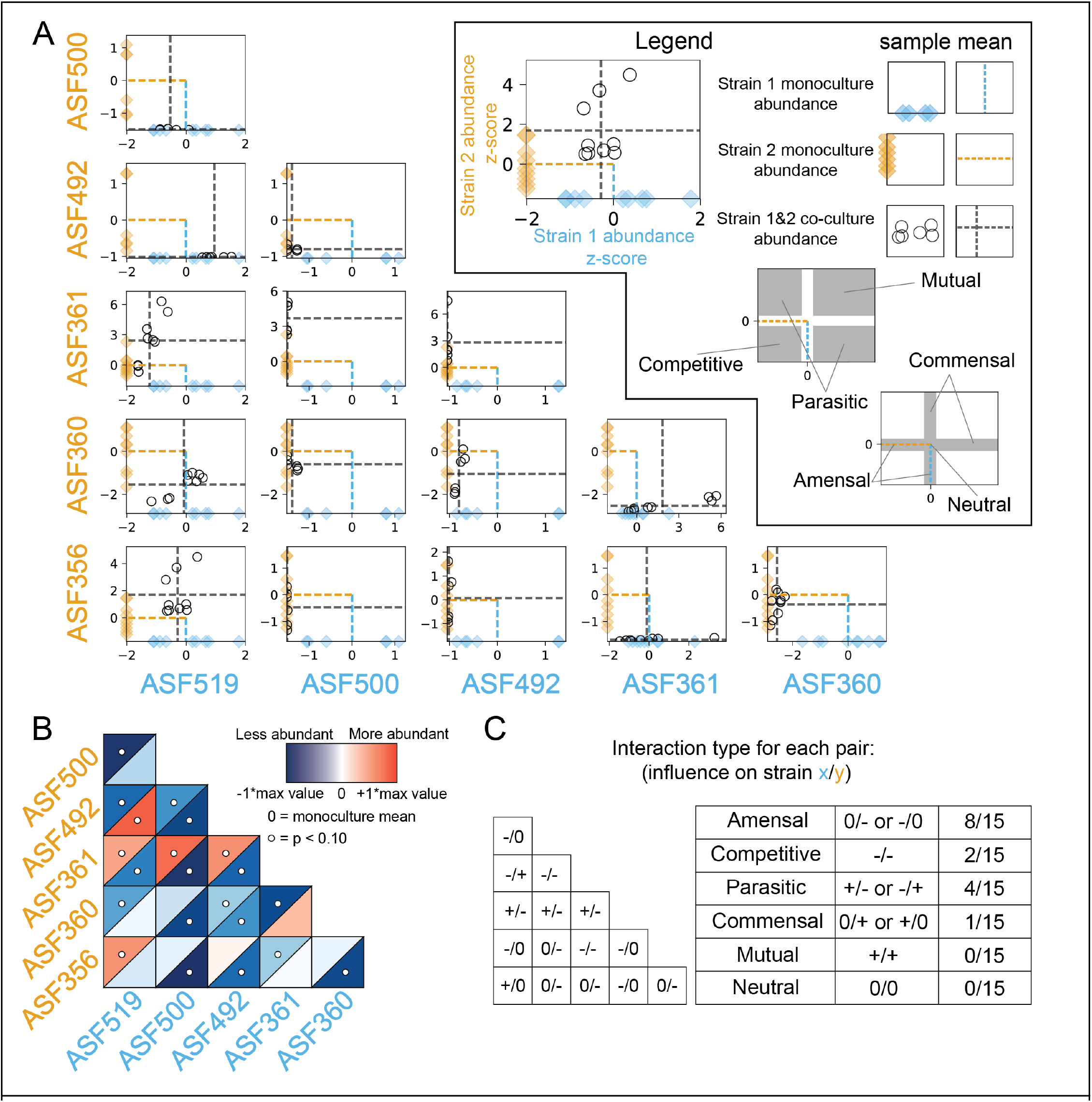
**A)** Abundance of each strain in monoculture and in co-culture determined via qPCR. In each subplot, the *x*-axis describes the abundance of the strain at the bottom of the column in sky blue text. Conversely, the *y*-axis describes the abundance of the strain labelled in orange text at the left of each row. The diamonds indicate the abundance of each strain in monoculture, with the mean shown by a dashed line of the same color. For each subplot, the abundance of each strain in the co-culture for the pair indicated by the row and column labels is shown by a black circle, with the mean co-culture abundances indicated by two grey dashed lines. Abundance values for each strain are z-score normalized using the mean monoculture abundance to center the data and the standard deviation of monoculture abundance to scale the data. N=9 for all samples except for monocultures and co-cultures containing ASF500 or ASF492, for which N=6. **B)** Heatmap of mean abundance of each strain in co-culture relative to abundance in monoculture. Blue indicates less abundant than monoculture, while red indicates more abundant than monoculture. The upper left triangle for each square describes abundance of the strain labelled on the left side of each row, while the lower triangle describes the abundance of the strain labelled on the bottom of each column. White circles indicate that the strain was differentially abundant when comparing mono-to co-culture (significance threshold of *p* < 0.10, Mann-Whitney U test with false discovery rate correction using Benjamini-Hochberg procedure). **C)** Summary of interspecies interactions. Non-zero interactions in the triangle indicate significant differential abundance as shown in Fig 2B.

### Metabolic repertoires within the altered Schaedler flora

To determine potential mechanisms governing the changes in growth observed in co-culture, we performed untargeted metabolomics on the spent supernatant from all samples in the growth experiments (using ^1^H NMR spectroscopy, see Materials and Methods). We updated and refined the metabolite peak annotations from experiments previously performed using the same medium and strains [21], resulting in 86 detected metabolites, 50 of which could be assigned an identity (compared to 36 of 85 metabolites previously assigned an identity). We identified several new metabolites involved in amino acid metabolism (serine, cystine, asparagine, glutamate, 2-oxoisocaproate, and isocaproate), nucleic acid metabolism (cytidine, cytosine, uridine monophosphate), and anaerobe-specific metabolism (isopropanol).

Based on the monoculture supernatant metabolomic profiles (Figure 3A), the ASF strains have fermentation repertoires similar to closely-related gut microbes. ASF360 and ASF361 both produced lactate, while ASF361 also produced acetate and formate. Other strains of *Lactobacillus intestinalis* and *Lactobacillus murinus*, the species that ASF360 and ASF361 are designated as, respectively, are generally identified as facultative heterofermentative lactic acid bacteria [22]. Heterofermentative lactic acid bacteria primarily ferment carbohydrates to lactate but may also produce additional acetate in some conditions. ASF356 produced the common fermentation end products acetate, propionate, succinate, and butyrate. Butyrate production is common in Clostridia that inhabit mammalian gastrointestinal tracts, and is often coupled with acetate production [23]. Propionate is the end product of three pathways identified in anaerobic organisms, of which the acrylate pathway and succinate pathway have been identified in *Clostridia spp*. [24]. ASF356 also produced isovalerate, isocaproate, and isobutyrate, which are common products of amino acid fermentation by some *Clostridia spp. [25]*. Butyrate and ethanol were the only common fermentation end products produced by ASF492. ASF492 has been proposed as the type strain for *Eubacterium plexicaudatum* [26], which was originally identified as producing butyrate and small amounts of acetate from glucose [27]. ASF500, a strain from the genus *Pseudoflavonifractor*, produced only formate and consumed less lactose than any other ASF strain. This suggests that lactose is not a preferred carbon source for ASF500 or another growth-limiting nutrient is only present at low abundance in the medium. ASF519 produced acetate, propionate, and succinate, consistent with previous reports on fermentation products of *Parabacteroides goldsteinii*, as well as formate [28]. ASF519 also produced many amino acids, including histidine, lysine, alanine, isoleucine, valine, proline, phenylalanine, glutamate, and methionine, suggesting ASF519 contains a comprehensive amino acid biosynthesis repertoire.

**Figure 3.**
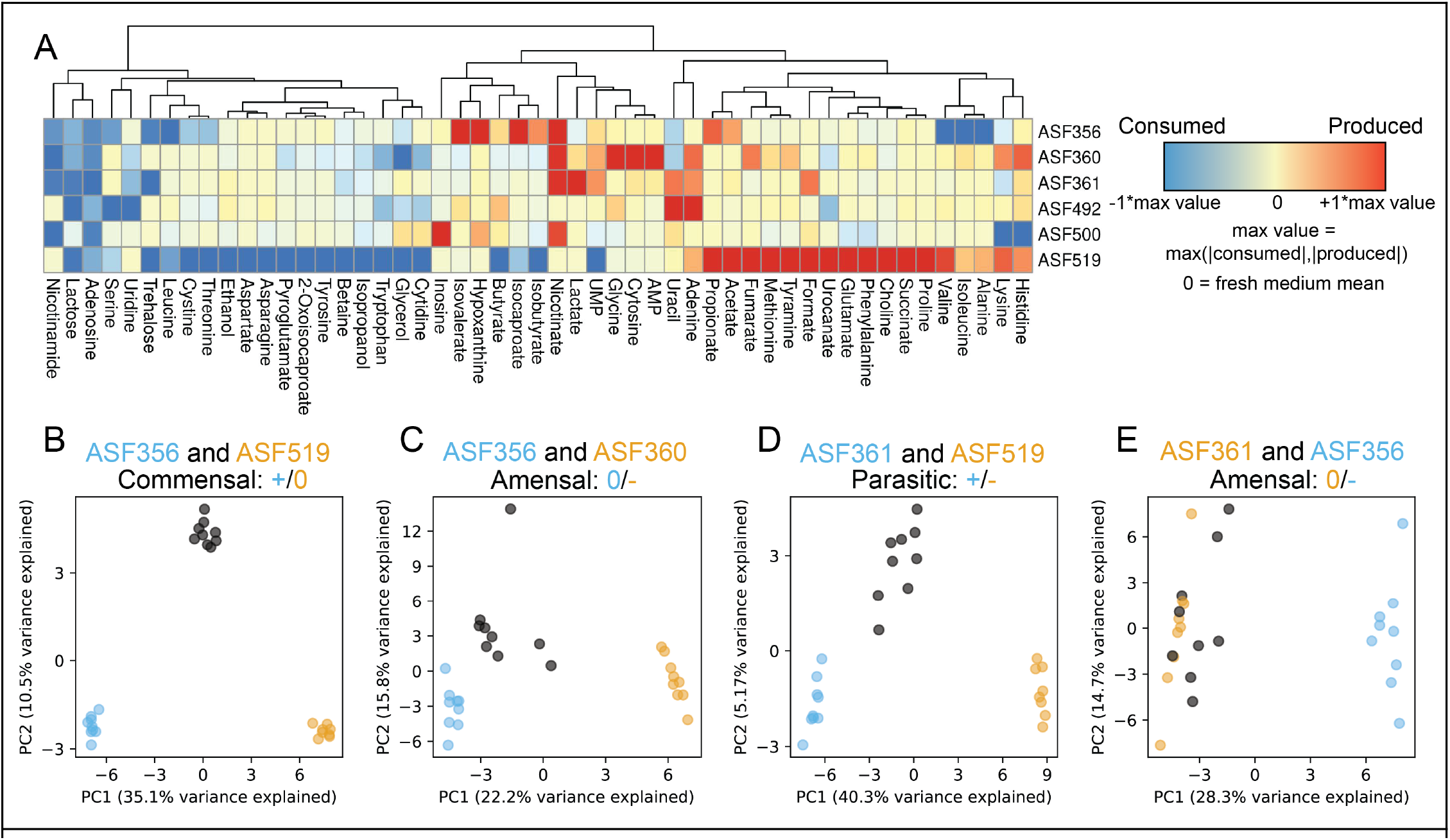
**A)** heatmap describing supernatant metabolomes for each strain after growth in monoculture. Red indicates higher concentration than fresh medium, while blue indicates lower concentration. Values are centered at 0 using the mean value from negative control replicates (N=9), then scaled between -1 and +1 using the maximum change in concentration (i.e. the max of the absolute values for the metabolite in all samples after centering using the negative control). Only metabolites for which an identity could be determined are shown. Hierarchical clustering of metabolites was performed using Euclidean distances and complete linkage. **B-E)** Principal component analysis (PCA) of monocultures and co-cultures from various strains. PCA was performed independently for each subplot. Within each subplot, sky blue circles correspond to monoculture supernatant metabolomes from the strain in sky blue text within the subplot title, and orange circles correspond to the strain indicated with orange text. Co-culture samples for the two strains in each subplot title are indicated by grey circles. Growth outcomes as described in Fig 2 are provided below in the title for each subplot.

### Co-culture leads to emergent metabolic behavior

Co-culture substantially altered the metabolome of pairings relative to each of the monoculture metabolomes for the strains involved. To detect and quantify the emergent metabolic behavior resulting from co-culture, we performed principal component analysis (PCA; see Materials and Methods) on the metabolic profile for pairs of strains. We performed PCA separately for each pair of strains, including samples from the monoculture for each strain in the pair and the co-culture samples for the pair. In cases where both strains grew in co-culture (i.e. no strong negative growth effect on one strain), the first principal component (PC1) separated monocultures by strain and the second principal component (PC2) separated monoculture samples from co-culture samples. This behavior was particularly strong in the case of ASF356 and ASF519 (Fig 3B). For ASF356 and ASF519, the loadings of PC2 suggest that co-culture increased production of propionate, glycine, and the amino acid fermentation products isovalerate, isocaproate, and isobutyrate, and increased consumption of multiple amino acids and lactose.

For pairings with a strong negative effect on one strain, the co-culture metabolomes were less similar to the negatively-affected strain than they were to the strain that did not experience the negative growth effect. For example, co-culture of ASF356 with ASF360 resulted in decreased growth of ASF360, and the co-culture samples are located closer to the ASF356 monoculture samples in PCA (Fig 3C). Although there is still an “emergent” co-culture effect observed in PC2 for this pair, the effect is also aligned with within-group variation (e.g. samples within each monoculture vary along PC2). The same trend is present for the metabolomes of ASF361, ASF519, and the co-culture of the two strains (Fig 3D). For strong negative growth outcomes (e.g. negative effect on ASF356 during co-culture with ASF361), the effect is more pronounced and there is less separation between monoculture and co-culture samples (Fig 3E).

### Co-culture enhances efficiency of resource utilization

Based on the metabolic differences between monoculture and co-culture samples identified via PCA, co-culture conditions substantially altered metabolic behavior. However, the mechanism that leads to this emergent metabolic behavior is unclear, and attempting to infer the mechanism may be confounded by changes in the abundance of each strain in co-culture. We sought to create a theoretical framework to infer metabolic interactions between strains in co-culture while controlling for changes in strain abundance. We provide two examples to motivate this framework (Fig 4A). In each example, we examine a single metabolite and the behavior of that metabolite in monoculture and co-culture for a pair of strains in which one strain experienced a growth benefit in co-culture and another experienced no change in growth in co-culture. Hypothetical metabolite 1 was produced by both strains, and a yield for that metabolite for each strain can be calculated by dividing its abundance by the abundance of each strain after growth in monoculture. In this case, the challenge is to determine whether the yield of that metabolite increased or decreased for either strain when the two strains are grown in co-culture. While the amount of a metabolite produced may increase in co-culture relative to in monoculture (e.g. short-chain fatty acids in co-culture of ASF356 and ASF519), a calculation that takes into account changes in strain abundance is necessary to determine whether the increase in metabolite abundance is truly emergent behavior rather than additive. Similarly, hypothetical metabolite 2 was produced by one strain and consumed by another in monoculture. Given the changes in strain abundance in co-culture, we ask whether the metabolite was cross-fed, and whether the metabolite might have contributed to changes in strain abundance observed in co-culture.

**Figure 4.**
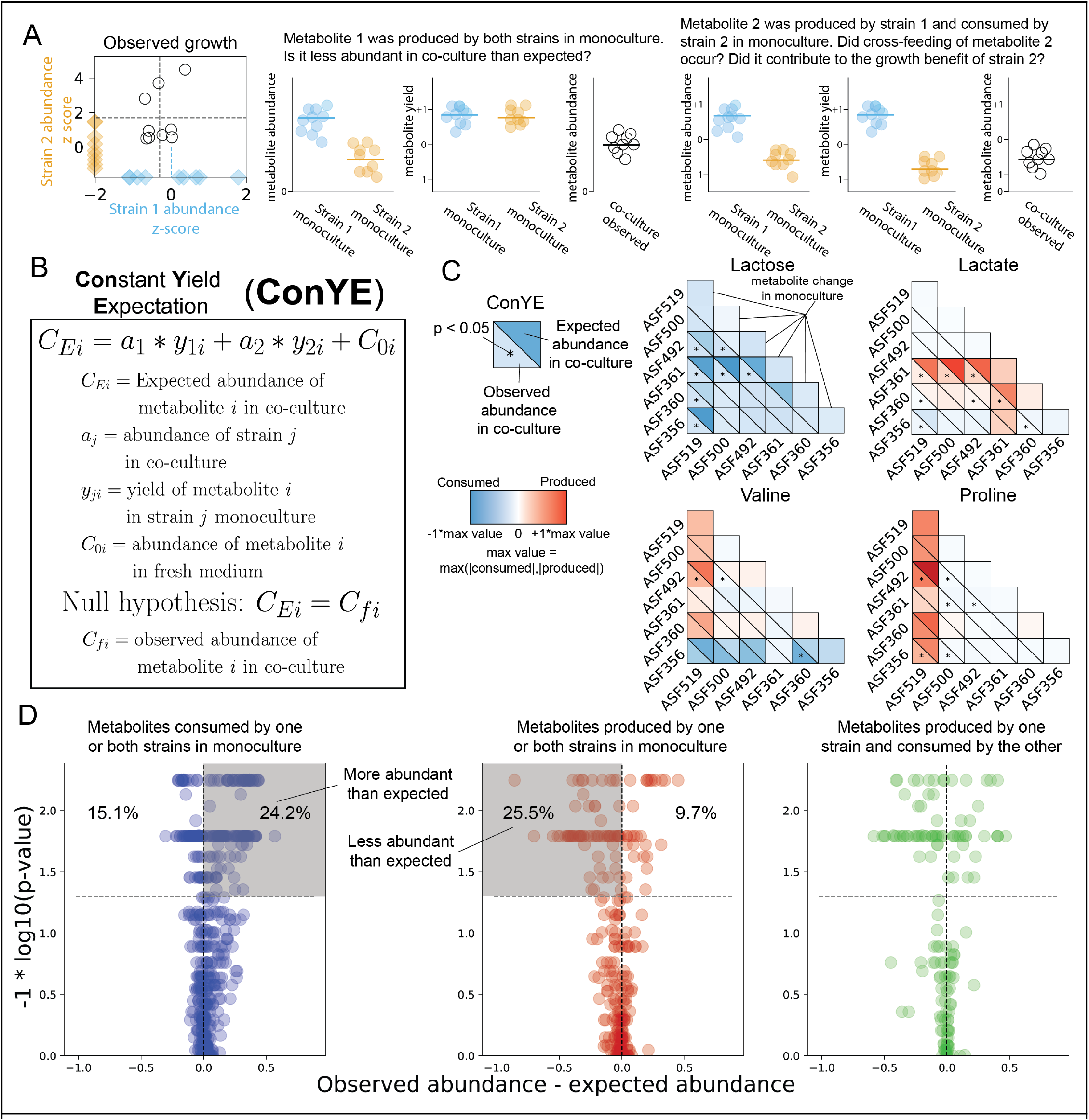
**A)** Hypothetical case representing examples where differential metabolite yield of a produced metabolite (metabolite 1) or cross-feeding (metabolite 2) may have occurred in co-culture given the growth outcomes for strain 1 and 2. **B)** Equation and procedure for the constant yield expectation (ConYE) model. **C)** Examples of ConYE results for Lactose, Lactate, Valine, and Proline. The diagonal represents monoculture behavior for each strain. Every other pair of triangles indicates the observed metabolite abundance in co-culture (lower left), the expected metabolite abundance in co-culture (upper right), and whether there was a significant difference between the observed and expected values. Max value for color scale determined by the max of the absolute value of metabolite z-scores across all monocultures and co-cultures. Null hypothesis testing was performed with Mann-Whitney *U*-Test with false discovery rate (FDR) control using the Benjamini-Hochberg procedure across all 1290 comparisons (15 co-cultures, each with 86 metabolites). Asterisk in lower left triangle indicates *p* < 0.05 for the given metabolite in the co-culture containing the indicated strains. **D)** ConYE results for all strain pairings for metabolites that were consumed by one or both strains in monoculture (left, blue), produced by one or both strains in monoculture (middle, red), or produced by one strain in monoculture and consumed by the other strain in monoculture (right, green). In each subplot, each point represents a single metabolite in a specific co-culture pair (i.e. up to 1290 points are represented across all subplots, representing all combinations of metabolites and unique strain pairs). The x axis describes difference between observed and expected metabolite abundance in co-culture (scaled as in panel C), and the y axis represents the FDR-adjusted p-value derived by comparing the ConYE model to observed metabolite abundances. Points above grey line have *p* < 0.05. Points in shaded region in left subplot (blue) are metabolites that were more abundant in co-culture than expected. Points in the shaded region in middle subplot (red) are metabolites that were less abundant than expected in co-culture. Percentages shown are the percentage of points in the labelled quadrant relative to the rest of the points in the subplot.

Based on the metabolic abundance profiles alone, it is unclear whether co-culture led to cross-feeding and/or resource competition, the two mechanisms of metabolic interaction that are based on resource allocation. If two strains consume a metabolite in monoculture, resource competition only affects growth of either strain if the metabolite is completely consumed or the concentration becomes low enough to affect growth indirectly (e.g. reduced transport/diffusion). In the case of cross-feeding, lower amounts of a metabolite produced by one strain could be due to reduced growth of that strain, rather than consumption of that metabolite by the other strain (Fig 4A). Inference of cross-fed metabolites or metabolites for which competition affected growth outcomes is thus complicated by changes in the total growth of individual strains in co-culture relative to monoculture. We developed the **Con**stant **Y**ield **E**xpectation (**ConYE**) model (see Materials and Methods) to account for variable growth and aid interpretation to identify metabolites for which consumption or production behavior changed in co-culture. Within the ConYE model, we assume each strain produces or consumes a fixed quantity of each metabolite per unit biomass (i.e. constant yield). We simulate expected metabolite quantities in co-culture by multiplying the mono-culture-derived metabolite yield for each strain by the observed abundance of that strain in co-culture, then summing the expected values for each strain and the initial quantity of the metabolite present in the fresh medium (Fig 4B). For each metabolite, we test the null hypothesis that the quantity of that metabolite in co-culture is equal to that predicted by the ConYE model. Rejecting the null hypothesis for a metabolite implies that co-culture caused at least one strain to alter metabolism of that metabolite relative to its own biomass production. Throughout Fig 4 and Fig 6, we illustrate the magnitude of deviation from expectation, as well as the associated p-value, for each metabolite in each co-culture condition using volcano plots (Fig 4D, all metabolites for all co-cultures shown).

We identified several patterns with the ConYE model results that were consistent across sets of many metabolites, for which representative examples are shown (Fig 4C). Metabolites that were commonly consumed in monoculture were often consumed less than expected in co-culture, especially when one strain in the co-culture experienced a growth benefit (e.g. lactose). For some strains, this pattern may arise because alternative metabolites are now available in co-culture that can be consumed to produce biomass, decreasing the amount of lactose required to produce a unit of biomass for that strain. Similarly, another pattern involves fermentation end products, which were generally less abundant than expected. For example, lactate, which was produced by ASF360 and ASF361, was less abundant than expected in 7/9 co-cultures containing either of the strains. Possible explanations for this pattern align with explanations for the first pattern; individual strains may be utilizing alternative metabolites to produce biomass, resulting in less production of their primary fermentation products. An alternative explanation is that other strains in the co-culture are consuming the fermentation end product, as may be the case for lactate (ASF356, ASF492, and ASF519 all consumed lactate present in the fresh medium). Similar explanations may fit the behavior of other metabolites that are not end products of fermentation, such as valine. Valine was consumed by some strains and produced by others, but the null hypothesis for valine was only rejected for 3/15 co-cultures. In cases where one strain produced a metabolite in monoculture (e.g. ASF519 producing valine) and another strain consumed the metabolite in monoculture (e.g. ASF356 consuming valine), failure to reject the null hypothesis even when one species experienced a growth benefit (e.g. ASF356 co-cultured with ASF519) suggests that a metabolite may have been cross-fed.

As demonstrated by considering these examples, interpretation of rejecting the null hypothesis can be informed by considering how the concentration of the metabolite changed in monoculture. If one or both strains consumed a metabolite in monoculture (Fig 4D, left, all co-cultures shown), rejecting the null hypothesis implies the metabolite was consumed more or less than expected, or that one of the strains produced the metabolite in co-culture (e.g. emergent production). Conversely, if one or both strains produced a metabolite in monoculture (Fig 4D, middle), rejecting the null hypothesis implies the metabolite was produced more or less than expected, or that one of the strains consumed the metabolite in co-culture (which, again, was not observed in monoculture for that strain). For both the production and consumption cases, cross-feeding is still possible, but in limited situations (e.g. emergent consumption or production by one strain).

In the case where a metabolite was consumed by one strain in monoculture and produced by the other strain in monoculture (Fig 4D, right), there are four possible interpretations if the null hypothesis is rejected. If the metabolite was less abundant than expected, then at least one of two conclusions is true: 1) the consumer metabolized more of the metabolite than expected, or 2) the producer produced less. If the metabolite was more abundant than expected, the opposite is true (producer produced more or consumer consumed less). If the null hypothesis is not rejected, the strains either maintained their production and consumption behavior from monoculture, or both scaled their consumption and production up or down, each in equal amounts. These interpretations, as well as their corresponding importance or relative contribution to a positive growth interaction for the consuming strain, are summarized in Fig 5.

**Figure 5.**
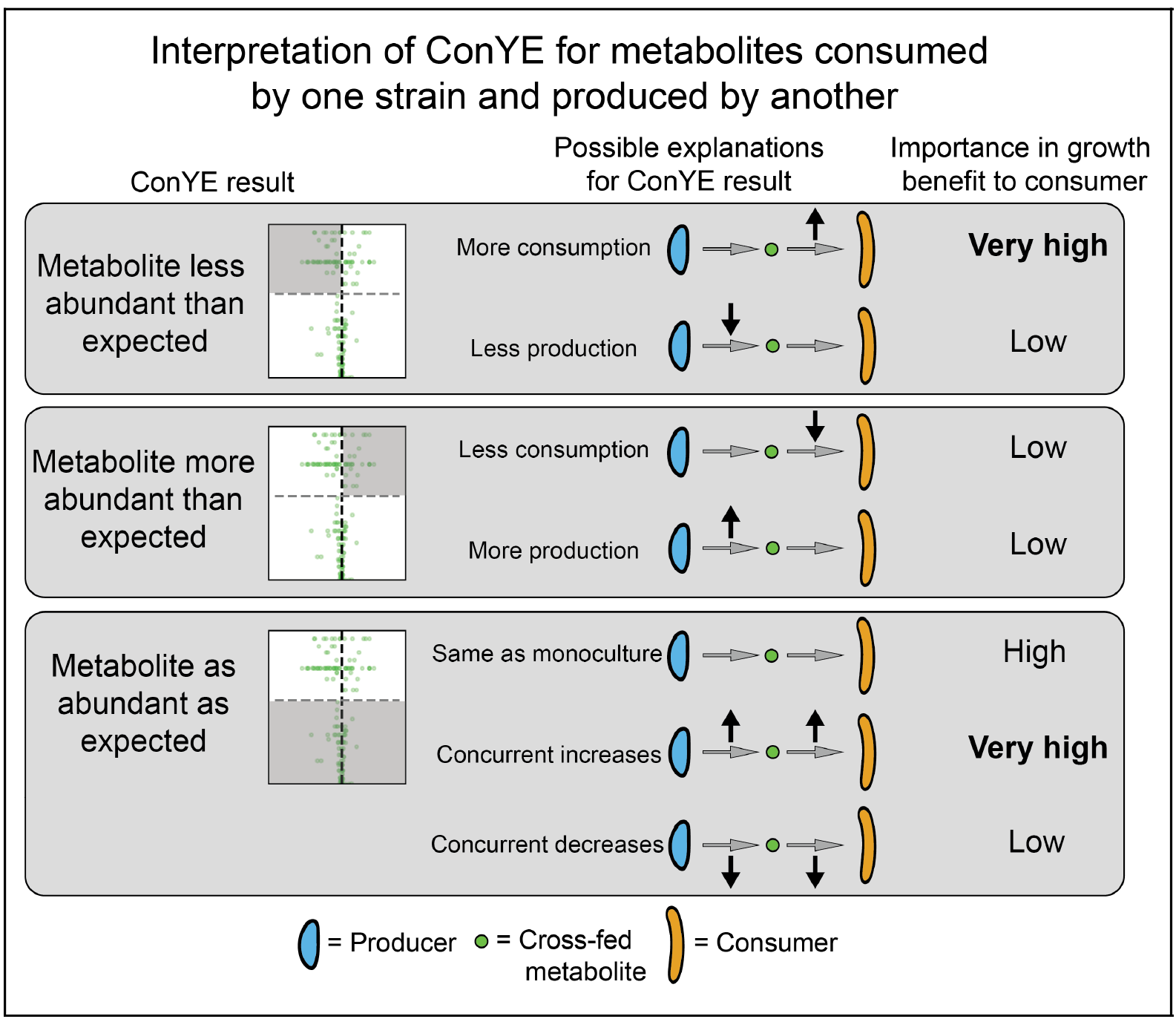
Possible interpretations of ConYE results for metabolites consumed by one strain in monoculture and produced by another strain in monoculture, as described in Results. In this scenario, the consumer experienced a growth benefit. The shaded region of each small volcano plot describes the points that fall into the category described to the left of the plot. In “Importance in growth benefit to consumer” column, the entry for each scenario assumes that consumption of the metabolite is coupled with biomass production.

### Niche expansion and cross-feeding occur with positive growth-modulating interactions

After applying ConYE to all co-cultures, the null hypothesis was rejected for 500/1290 metabolites (38.8%), suggesting that co-culture alters metabolism of a substantial portion of metabolites when taking into account changes in growth that occur during co-culture. For metabolites that were consumed by one or both strains in monoculture, the amount consumed per unit of strain growth generally decreased in co-culture if the null hypothesis was rejected. Specifically, of the 624 instances of metabolites that fell into this category, 151 (24.2%) were significantly more abundant than expected in co-culture, whereas 94 (15.1%) were less abundant than expected. Of the 278 instances of a metabolite being produced by one or both strains in a pairing in monoculture, 71 (25.5%) were less abundant than expected, while 27 (9.7%) were more abundant than expected. Thus, although co-culture often resulted in a greater quantity of a metabolite being produced relative to either monoculture (i.e. metabolites driving monoculture and co-culture separation in PCA, Fig 3B), the amount produced relative to growth of each strain decreased for most metabolites. Similarly, the amount of each metabolite consumed relative to biomass in co-culture generally decreased. These results suggest that these co-cultures can improve overall biomass production (as in Fig 1C) through niche expansion (e.g. consuming metabolites they did not consume in monoculture) and/or cross-feeding rather than increasing consumption of metabolites they did not fully deplete in monoculture. Indeed, 90/1290 (6.98%) metabolites were not consumed or produced by either strain in monoculture, yet were consumed when the two strains were in co-culture.

These distinct ConYE trends are enriched in cases with positive growth interactions (Fig 6A). When considering only pairings with a positive growth effect for at least one strain, there were 219 metabolites that were consumed by one or both strains in monoculture. Of these 219 metabolites, 138 (63.0%) were more abundant than expected, while only 6 (2.74%) were less abundant than expected. Of the 88 metabolites produced by one or both strains in monoculture for these co-cultures, 51 (58.0%) were less abundant than expected, and only 5 (5.68%) were more abundant than expected. Taken together, these results indicate that co-cultures with positive interactions are able to more efficiently utilize resources than co-cultures without positive interactions or monocultures.

**Figure 6.**
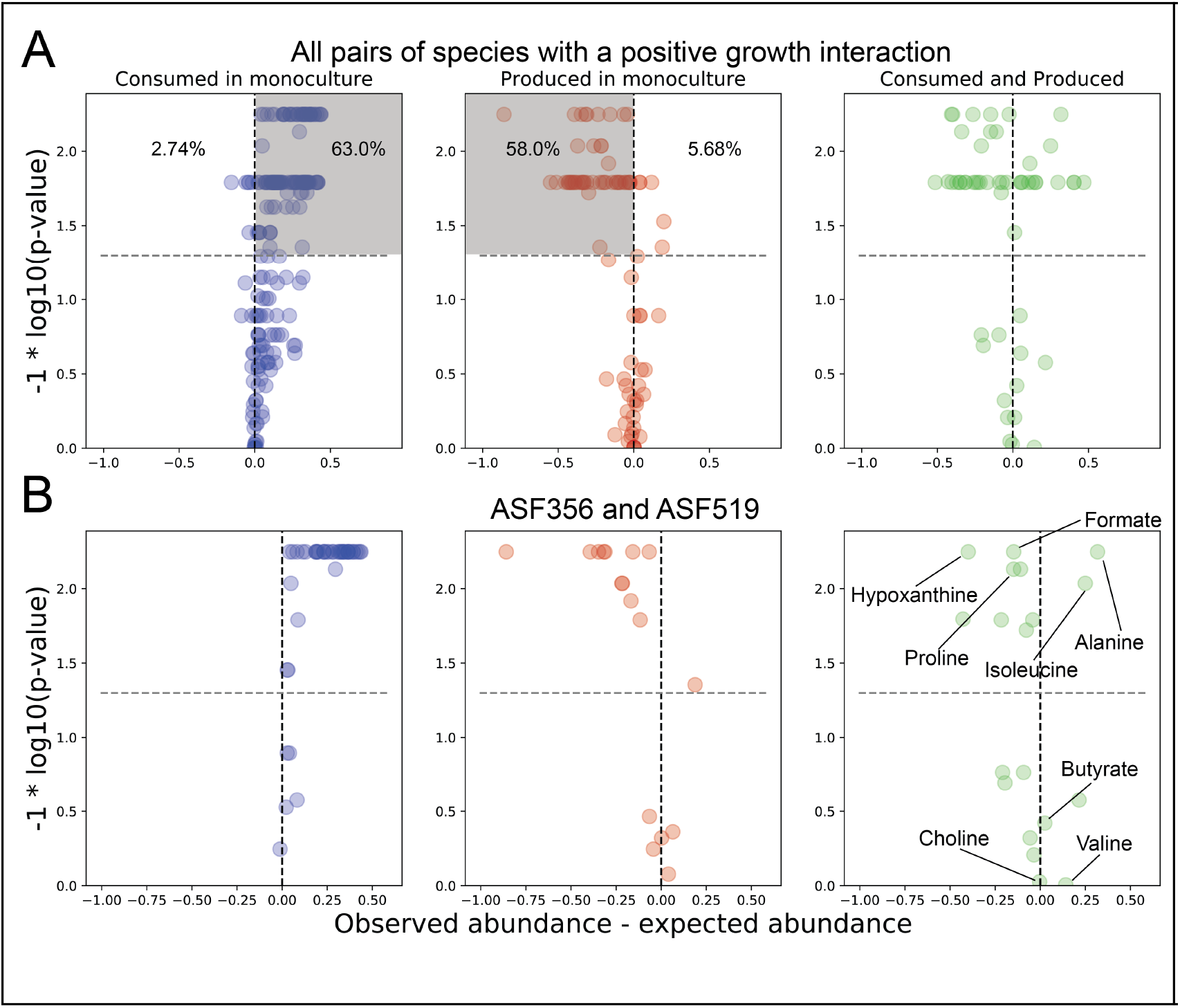
**A)** ConYE results for all metabolites from co-cultures in which at least one strain experienced a positive growth interaction. Shaded quadrants represent consumed metabolites that were more abundant than expected (left) or more abundant than expected (middle). Percentages shown represent that number of metabolites within the volcano plot that fall in the quadrant. **B)** ConYE results for the co-culture containing ASF356 and ASF519. Metabolites on right for which p > 0.05 are labeled unless they could not be assigned an identity. Metabolites for which p < 0.05 are labeled if assigned an identity and abs(x) > 0.10 for that metabolite. X and y axes are scaled as in Fig 4.

There are two mechanisms that may enable this: niche expansion (consumption of metabolites not consumed in monoculture) and cross-feeding. In co-culture, the subset of strain pairs with positive interactions consumed 30 metabolites that were not consumed by either species in monoculture. Interestingly, all 30 instances of emergent metabolite consumption were carried out by ASF361+ASF492, ASF361+ASF500, and ASF492+ASF519, while the remaining two pairs (ASF356+ASF519 and ASF361+ASF519) had 0 cases of emergent consumption. Given this result, it is likely that the growth benefits that occurred for ASF356+ASF519 and ASF361+ASF519 are due to cross-feeding, while the growth benefits for the other positive interaction pairs are at least in part due to niche expansion.

### Identifying cross-fed metabolites and evaluating feasibility *in silico*

We next sought to investigate potential cross-fed metabolites from ConYE for co-cultures with positive growth interactions in order to find a mechanism that explained, at least in part, the growth benefit. For this task, we focused on the co-culture of ASF356 and ASF519 to exclude co-cultures which may have engaged in niche expansion (and therefore cross-feeding may have played a small role in observed growth benefits) and to remove the need to consider additional confounding factors introduced by a strong negative growth interaction (e.g. negative impact on ASF519 growth in co-culture with ASF356). For the co-culture of ASF356 and ASF519, there were 7 named metabolites that were consumed by ASF356 in monoculture and produced by ASF519 in monoculture (Fig 6B). Of those 7 metabolites, tyramine, valine, and choline did not result in rejecting the null hypothesis (e.g. they were as abundant as expected). Isoleucine and alanine were more abundant than expected, and proline and formate were less abundant than expected. Isoleucine and alanine may have been cross-fed, but given that they were more abundant than expected, consumption of these metabolites only contributed to enhanced growth if ASF519 also produced less of these metabolites than expected (as in middle panel of Fig 5, where ASF519 is the producer and ASF356 is the consumer). Proline and formate were both less abundant than expected, so were either consumed by ASF356 more in co-culture than in monoculture (and thereby cross-fed) or produced less by ASF519 in co-culture than in monoculture (as in top panel of Fig 5).

ConYE can identify metabolites that are potentially cross-fed, but the actual behavior of each strain in co-culture with respect to that metabolite is difficult to infer using existing experimental techniques. Because we can only evaluate the co-culture behavior based on an expectation derived from monoculture behavior, it is still possible that co-culture leads to reduced production and consumption of those metabolites rather than cross-feeding. We sought to provide orthogonal evidence for ConYE results by evaluating the potential for metabolites to increase the growth rate of a strain in monoculture, reasoning that ConYE may produce false-positive inferences if metabolites are not actually coupled with biomass production. We chose to support inferences made using ConYE by building and applying Genome-scale metabolic network reconstructions (GENREs). GENREs are mathematical representations of all metabolic reactions that an organism can carry out, and have been used extensively to predict the effect of environmental conditions on growth of bacterial species [29]. We created an ensemble of 100 GENREs for each strain in this study to gain greater confidence in cross-feeding predictions and to enable predictive modeling of metabolism in future studies (Fig 7A and 7B; See Materials and Methods). For each metabolite, we evaluated its impact on growth of individual strains by performing ensemble flux balance analysis (EnsembleFBA) [30] to predict the growth rate of the strain without the metabolite available and with the metabolite available in excess (Materials and Methods). We performed this procedure for the candidate metabolites involved in cross-feeding interactions that increased growth of ASF356 when co-cultured with ASF519 (in which case ASF356 experienced a growth benefit). If a metabolite increases the predicted *in silico* growth rate when available in excess, we take that as parallel evidence to support or oppose the ConYE-based inferences. EnsembleFBA results are summarized in Fig 7C for all metabolites except tyramine, which was not present in any GENREs within the ensemble for ASF356.

**Figure 7.**
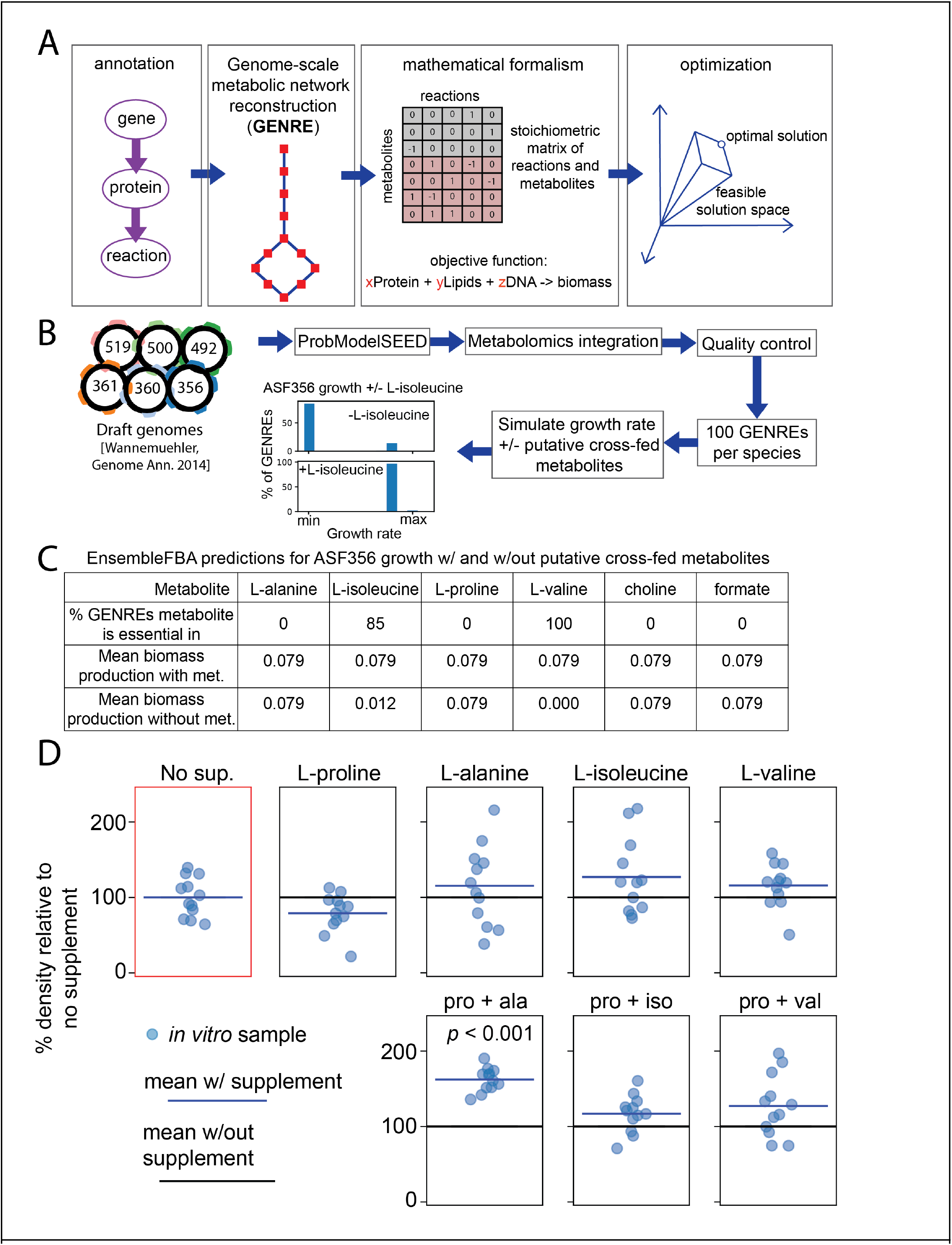
**A)** Process for generating and applying a genome-scale metabolic network reconstruction (GENRE). **B)** Novel pipeline for constructing and analyzing ensembles of GENREs for the ASF strains. Details for each step in Materials and Methods. **C)** EnsembleFBA predictions of the influence of potentially cross-fed metabolites on growth of ASF356. Biomass production values are for flux through the biomass reaction in units of hour^-1^, and a metabolite was considered essential if removal led to flux through the biomass reaction of less than < 1 E-5/hour. **D)** OD600 of ASF356 monocultures after 72 hours of growth in supplemented media conditions.”No sup.” had no supplement added, while conditions with a single amino acid were supplemented at 1.25g/L. In conditions with multiple amino acids (bottom row), each of the two amino acid are still supplemented at 1.25g/L. “Pro + ala”, “pro + iso”, and “pro + val” conditions include L-proline with L-alanine, L-isoleucine, or L-valine, respectively.

Valine was essential for growth of ASF356 in all 100 GENREs in its ensemble. Isoleucine was essential for 85/100 GENREs but had no effect on growth for the other 15 GENREs, while absence of the rest of the potentially cross-fed metabolites had no effect on predicted growth rate. Given the *in silico* essentiality of valine and the ConYE results indicating valine was as abundant as expected, valine may have conferred a growth benefit to ASF356 if cross-fed between ASF356 and ASF519. While isoleucine was essential for growth of the majority of GENREs in the ASF356 ensemble, there was a subset of GENREs in which its removal had no effect. Given this *in silico* uncertainty, as well as the ConYE result which indicated it was more abundant than expected in co-culture, suggest that cross-feeding of isoleucine may not have influenced growth of ASF356 as much as valine. Alanine, proline, choline, and formate were not essential and did not influence predicted growth rates *in silico*. Critically, however, this analysis indicated that availability of any of the individual metabolites in excess did not confer a growth benefit relative to the unsupplemented medium.

### Validation of an inferred cross-feeding interaction

Given the lack of an *in silico* prediction for supplementation of any metabolite to increase the growth rate for ASF356, we considered mechanisms through which the metabolites discussed above may interact with each other to influence growth, rather than acting in isolation as we considered thus far. ASF356 belongs to the *Clostridium* genus, throughout which amino acid fermentation via the Stickland reaction is common [25]. The Stickland reaction involves coupling the oxidative deamination of one amino acid with the reductive decarboxylation of another amino acid, producing two short-chain fatty acids or branched chain fatty acids that each contain one fewer carbon than the respective amino acid from which they were derived [31]. Proline, glycine, hydroxyproline, and ornithine are strong Stickland reaction electron acceptors, while alanine, valine, leucine, and isoleucine are strong electron donors. We observed that ASF356 consumed proline, a strong electron acceptor, and all the listed electron donors in monoculture, while ASF519 produced proline, alanine, valine, and isoleucine and consumed leucine in monoculture. In co-culture, ConYE indicated that proline was significantly less abundant than expected, suggesting it was consumed more per unit biomass in co-culture than in monoculture. Given this observation and the lack of growth rate increase predicted *in silico* with excess proline available, we hypothesized that proline was of critical importance to the growth benefit for ASF356 in co-culture with ASF519, but depended on the presence of suitable electron donors. Behavior varied amongst the electron donors that may pair with proline in the Stickland reaction: isoleucine and alanine were more abundant than expected, while valine was as abundant as expected. The Stickland fermentation product for proline is 5-aminovalerate, which we could not identify within our NMR spectra due to spectral overlap with other metabolites and lack of signal in regions unique to 5-aminovalerate. The products for isoleucine, valine, and alanine are valeric acid (not detected), isobutyrate (as abundant as expected), and acetate (less abundant than expected), respectively. Decreased abundance of leucine, which is fermented to isovalerate (less abundant than expected), in co-culture suggests decreased consumption by ASF356 or increased consumption of isovalerate by ASF519, which consumed isovalerate in monoculture.

To test the hypothesis that ASF356 experiences a growth benefit in the presence of proline and suitable electron donors, we grew ASF356 in media supplemented with proline, alanine, isoleucine, valine, or each combination of the three electron donors (alanine, isoleucine, valine) with proline. Although NMR spectroscopy cannot differentiate between amino acid isomers, we assumed all amino acids consumed and produced were the L- isoform. This assumption is unlikely to to impact our results, since tryptone, the major protein source in the medium used, is a casein digest which contains only L- isoforms. Additionally, organisms conducting Stickland fermentation of proline generally possess a proline racemase, since D-proline is the isoform that is fermented [32]. We did not use leucine in these experiments because it was consumed by both ASF356 and ASF519 in monoculture, thus was unlikely to be cross-fed in co-culture. Supplemented conditions contained either 1.25g/L of a single amino acid or 1.25g/L of each Stickland pair (e.g. 1.25g/L proline and 1.25 g/L alanine). Only the monoculture supplemented with both proline and alanine was significantly denser than monoculture with no supplement (Fig 6D, P < 0.05, Mann-Whitney *U*-test with false discovery rate control using Benjamini-Hochberg procedure), suggesting that Stickland fermentation of proline and alanine contributes to growth of ASF356. Given that the ConYE results indicated that alanine was more abundant in co-culture than expected, the results of the supplementation experiment imply that production of alanine by ASF519 was likely reduced in co-culture with ASF356, or that alanine was used more efficiently by ASF356 in co-culture than in monoculture. Additionally, the lack of growth benefit conferred by concurrent supplementation of proline with isoleucine or valine suggests the growth benefit attributable to pairing either electron donor with proline either did not occur or the effect is too small to detect given our sample size. Formate can also be used as an electron donor for proline reduction as an alternative to the conventional Stickland pairs [33], which we did not take into consideration when designing these experiments. Formate was produced by ASF519 and consumed by ASF356 in monoculture, and was less abundant than expected in co-culture according to ConYE. Thus, formate may have also contributed to the growth benefit of ASF356 in co-culture with ASF519.

We also tested the effect of concurrent supplementation with Stickland pairs *in silico*, however the GENREs do not contain Stickland fermentation reactions and thus the predictions were no different than in the single amino acid supplement cases. Stickland reactions (e.g. D-proline reductase and glycine reductase) are absent from reaction databases used to construct and curate GENREs such as the ModelSEED biochemistry database [34] used in this study and BiGG models [35], but are present in the AGORA resource of semi-automatically generated GENREs for gut microbes [36]. None of the genes involved in Stickland fermentation have been identified in the genome of ASF356 as of this writing (determined via searching the annotated genome for ASF356 in the PATRIC database [37] and targeted BLAST against D-proline reductase gene subunits).

## Discussion

In this study, we used data-driven methods to identify metabolic signatures that may contribute growth modulation in bacterial co-cultures, proposed mechanisms by which a specific signature may arise, and verified that growth of the benefiting strain can be enhanced via this mechanism. The mechanism we verified experimentally, Stickland fermentation of proline and alanine, is widely distributed in proteolytic *Clostridia [25, 31]* including pathogenic species such as *Clostridium difficile* [38]. While the ability of species inhabiting the mammalian gut to perform Stickland fermentation is widely studied, this study is the first to our knowledge to connect Stickland fermentation to a specific, bidirectional interspecies interaction that modulates growth. Given that some of the end products of Stickland fermentation were present at low concentrations in the fresh medium, and that ASF519 consumed them in monoculture (isobutyrate, isovalerate, and isocaproate), our data suggest that this interaction may be mutually beneficial and bidirectional. This observation supports some theoretical motifs for metabolism in the gastrointestinal tract proposed in the literature, such as the model of carbon and nitrogen flow proposed by Fischbach and Sonnenburg in which *Clostridia* (e.g. ASF356) ferment amino acids, providing ammonium and other amino acid fermentation products to *Bacteroidetes* (e.g. ASF519) [8]. Indeed, ASF356 and ASF519 are highly co-located along the mouse gastrointestinal tract, however the relevance of this observation is unclear given a microbiota as restricted in size as the ASF [39]. This kind of interaction has direct relevance to enteric pathogens, as emerging evidence indicates that *in vivo* utilization of proline via Stickland fermentation is highly active in *Clostridium difficile* during sustained infection in mice [40, 41].

We expect that the growth outcomes observed in the co-culture of ASF356 and ASF519 are due to a multitude of interactions that each have a small effect. An alternative mechanism by which co-culture could enhance growth is consumption of growth-inhibiting products. Although we did not explore them in this study, ConYE identified several cases in which this may have occurred. For example, in the co-culture of ASF361 and ASF519, lactate, hypoxanthine, AMP, and UMP were all metabolites produced by ASF361 and consumed by ASF519 that were less abundant than expected in co-culture. Of these metabolites, lactate is known to have potent antimicrobial properties [42], thus is a reasonable candidate for this mechanism.

We developed ConYE to make sense of growth-modulating interactions within this study, but the framework can easily be extended to study interspecies interactions that alter other phenotypes of interest. For example, the same analyses conducted here could be performed using consumption of a substrate of interest, such as lactose, to calculate metabolite yields as a function of that substrate rather than as a function of strain abundance. In this scenario, ConYE could be used to identify co-culture pairings that enhance conversion of lactose to a metabolite of interest, and identify cross-fed metabolites that contribute to enhanced yield of that metabolite of interest.

There is increasing interest in developing methods for inference of interactions between microbes from various data types and environments [10, 43, 44]. These methods have primarily been focused on discovering interspecies interactions and the role they might play in ecosystem function, rather than ascribing mechanism to those interactions. However, interspecies interactions are likely to be highly context-dependent, so more detailed knowledge about mechanisms of interaction is necessary to generalize these findings [45]. Several approaches that integrate the metabolic and spatial environment have been developed that account for context dependency [46-48], but limitations in biochemical knowledge across the bacterial tree of life limit their broad application to other organisms. Similar approaches have been applied to simplified versions of complex communities such as the human gut microbiota [48-50]. However, the poor experimental tractability of these systems makes testing predicted interspecies interactions challenging and thus they are left unvalidated.

We have developed a novel experimental and computational pipeline to probe interspecies interactions and infer feasible metabolic mechanisms of interaction, generating testable hypotheses. Understanding mechanisms of interspecies interaction and the environmental conditions that induce them is a prerequisite to engineering communities with specific therapeutic or industrial value. For generalizable methods for predicting interspecies interaction using mechanistic models to be successful, methods must be validated. This task will require a substantially larger set of observed interspecies interactions than are presented here, or are available in the literature, from which to derive generalizable principles. Extending our approach and similar methods to defined communities across conditions that are more diverse, both in terms of resource availability and spatial structure, will begin to make predictive modeling of interspecies interactions tractable.

## Materials and Methods

### Strain maintenance

All strains are identified within the manuscript by the original isolate designation numbers for the ASF [12]. ASF457 was excluded from the study due to lack of detectable growth in the experimental medium, and ASF502 was excluded due to inconsistent growth in the experimental medium. Stock vials for all ASF strains were maintained at -80°C in 50% glycerol, 50% brain-heart infusion (BHI) medium (see media formulations for the composition of brain-heart infusion media used in this study). All strains were grown in an anaerobic chamber (Shellab BACTRONEZ, Sheldon Manufacturing, Inc., Cornelius, Oregon, USA) filled with mixed anaerobic gas (5% CO_2_, 5% H_2_, 90% N_2_). Anaerobic conditions were ensured through the use of palladium catalysts (baked at 120°C when outside the chamber and rotated daily when first entering the chamber) and anaerobic indicator strips (Oxoid, Basingstoke, UK).

### Media formulation

Supplemented Brain–Heart Infusion medium, referred to as BHI throughout the manuscript: Brain–Heart Infusion base (37g/L, BD, Franklin Lakes, NJ, USA), supplemented with yeast extract (5 g/L), 0.2mL of vitamin K1 solution (0.5% vitamin K1 dissolved in 99.5% ethanol), 0.5mL/L of hemin solution (0.5 g/L dissolved in 1% NaOH, 99% deionized water), L-cysteine (0.5 g/L) and 5mL/L each of newborn calf serum, horse serum, and sheep serum. Vitamin K1, hemin, and all sera were added after autoclaving the medium. For preparation of agar plates, agar was supplemented at 12g/L.

Modified Lennox LB medium, referred to as mLB throughout the manuscript: 30g/L LB base in powder form (Sigma, St Louis, MO, USA) was combined with 0.376g/L L-cysteine (Sigma), 39mL of a mineral salts solution (containing 6g/L KH_2_PO_4_, 6g/L (NH_4_)_2_SO4, 12g/L NaCl, 2.5g/L MgSO_4_•7H_2_O, 1.6g/L CaCl_2_•2H_2_O, all dissolved in deionized water), 15mL/L of hemin solution (0.5 g/L dissolved in 1% NaOH, 99% deionized water), 0.3mL of vitamin K1 solution (0.5% vitamin K1 dissolved in 99.5% ethanol), 15mL/L of lactose solution (5g/L lactose dissolved in deionized water) and 15mL/L of tween 20 solution (1g/L tween 20 dissolved in deionized water). All supplements made using deionized water, or that could not be autoclaved, were filter-sterilized using a 0.22μm membrane (except the sera).

#### *In vitro* monoculture and co-culture growth experiments in 12-well plates

Strains were inoculated from frozen stock to grow a dense lawn on agar plates containing BHI media. Prior to inoculation, all agar plates were equilibrated inside an anaerobic chamber for at least 24 hours. Inoculated plates were incubated for 3-9 days before being used to start overnight cultures. For overnight cultures, 50mL of mLB broth in a 500mL glass flask was inoculated using a generous streak from the lawn of each strain, then each flask was covered with a Breathe-Easy membrane (Diversified Biotech, Dedham, Massachusetts, USA). After 18-24 hours of incubation at 37°C, overnight cultures were transferred to 50mL conical tubes, sealed, transferred out of the chamber, and centrifuged at 1500 xg for 5 minutes. After centrifugation, samples were transferred into the chamber, supernatant was poured off, and pellets were resuspended in 12.7mL mLB broth. The resuspension for each species was then diluted to make inoculant with the same concentration of cells as a solution at an optical density of 0.01, measured at OD600 with 100uL of sample volume in a flat-bottom 96-well plate (with a well diameter of 0.64cm). This final inoculant was then used to inoculate mLB broth in 12-well plates. For monoculture samples, 100uL of inoculant was added to 2.9mL of media. For co-culture samples, 100uL of each strain’s inoculant was added to 2.8mL of media. 12-well plates were covered with a Breathe-Easy membrane, then the 12-well plate lid was placed on top of the membrane. Inoculated, covered 12-well plates were incubated at 37°C for 72 hours.

After 72 hours of incubation, 12-well plates were removed from the incubator and membranes were opened for each well using a razor. For each well, the sample was mixed by pipetting 900μL three times, then 1.8 mL of sample was transferred to a 2mL snap-cap tube. 200μL of sample was also collected after mixing and transferred to a 96-well plate to measure optical density at OD600. Samples within 2mL tubes were then transferred out of the chamber and centrifuged at 18407 xg for 2 minutes. After centrifugation, supernatant was poured directly into a 3mL syringe attached to a syringe pump filter (0.22μm pore size, mixed cellulose ester filter) and filtered into a 2mL snap-cap tube. Cell pellets were then resuspended in 400uL Qiagen lysis buffer (Buffer ASL, Qiagen) and vortexed until thoroughly mixed. Resuspended pellets and spent media were then frozen at -80°C.

To ensure reproducibility, 3 experiments were performed in which independent overnight starter cultures were used to inoculate 3 samples per monoculture and co-culture condition (resulting in 9 total replicates). For the third experiment, ASF492 and ASF500 did not appear to grow in monoculture according to both OD600 and qPCR, and their metabolomes did not appear significantly different than any negative controls, so all sample groups containing ASF492 and ASF500 in the third experiment were excluded from analyses. As a result, those sample groups have N = 6 replicates throughout the study.

#### *In vitro* amino acid supplementation experiments

Inoculant for ASF356 was prepared as described for monoculture and co-culture experiments in 12-well plates. Solutions of amino acids in deionized water were filter-sterilized (0.22μm pore size) and transferred to the anaerobic chamber and allowed to equilibrate for one week. Equilibrated solutions were mixed with liquid mLB broth (prepared as described previously), generating solutions that contained 90% mLB and 10% supplement by volume. Final concentrations were 1.25g/L for single amino acid supplements and 1.25g/L of each of two amino acids for supplements containing two amino acids (i.e. individual amino acids are at the same concentration in supplements containing one or two amino acids). 96-well plates were filled with 193μL media and 7μL inoculant (approximately the same initial density as in 12-well experiments), covered with a Breathe-Easy membrane, then incubated at 37°C for 72 hours. After incubation, the 96-well plates were removed from the anaerobic chamber, the breatheasy membrane was peeled off each plate, and the OD600 was measured in the 96 well plate.

### DNA extraction

Zirconia beads (1mm; BioSpec Products) were added to samples in ASL buffer (QIAmp Stool kit), and samples disrupted using a Mini-Beadbeater (15s, two times), followed by heat treatment (5 min, 90°C, 800rpm; Eppendorf Thermomixer). Debris was pelleted (14,000 xg, 1 min), and 400μL transferred to a QIAcube rotor adapter. Total DNA from each sample isolated on the QIAcube using the ‘human stool’ protocol provided by the manufacturer and stored at -20°C prior to PCR. Purified DNA used for standards in PCR assays was quantified using the DeNovix dsDNA kit. DNA standards were adjusted to 2ng/μL and diluted 10-fold to generate standard curves in PCR assays.

### Hydrolysis probe-based qPCR assay design

4-plex (including ASF356, ASF492, ASF502, and ASF519) and 3-plex (ASF360, ASF361, and ASF500) hydrolysis probe-based quantitative polymerase chain reaction (qPCR) assays were designed to quantify the abundance of each strain’s DNA with high specificity and throughput. Probe and primer design began with the *groEL* gene, which encodes the highly-conserved molecular chaperone protein GroEL, as a putative target. The National Center for Biotechnology Information PrimerBLAST web interface was used to identify PCR targets for each strain with minimal sequence similarity with any region in another strain’s genome [51]. PCR products ranging from 70-200 base pairs with a calculated melting temperature between 57°C and 63°C were determined, requiring at least two mismatches with unintended targets, with at least two mismatches occurring within the last five base pairs at the 3’ end. We screened the top three primer pairs for each strain returned by PrimerBLAST for sensitivity and specificity using standard SYBR green chemistry, and determined that all primers for ASF360, ASF361, and ASF500 had poor sensitivity. To identify alternative PCR products for ASF360, ASF361, and ASF500, we performed BLAST for each putative gene in each strain against all other putative genes in ASF strains. For genes with no hits (E-value > 1.0 for all comparisons), we attempted primer design using Primer-BLAST until a gene was found for each strain with at least 4 suitable primer pairs. All 4 primer pairs for the remaining ASF strains were screened for specificity and sensitivity and at least one suitable primer pair was found for each strain.

For all 7 strains in the study, probes were then designed for each primer pair. The 7 strains were split into a 3-plex and 4-plex reaction based on typical density observed experimentally, with strains growing to higher densities in the 4-plex reaction and strains growing to lower densities in the 3-plex reaction. For each probe in each reaction, we performed multiple sequence alignment using Clustal Omega [52]. Suitable probe sequences were identified manually according to five criteria: 1) maximize the number of mismatches at the 5’ end of the probe, 2) probe length between 20-30 base pairs, 3) estimated melting temperature around 66-70°C, 4) 35-65% GC content, and 5) no G or C at the 5’ end of the probe. Final primers, products, probe sequences, and accompanying probe fluorophores and quenchers are provided in Supplemental Table S1.

Primers and probes were synthesized by Integrated DNA Technologies, Inc. (Coralville, Iowa, USA). PerfeCTa 5X MultiPlex qPCR ToughMix (Quantabio, Beverly, MA) was used for all reactions. Each PCR reaction (20μL total volume) contained ToughMix (1X concentration), 300nM of each forward primer, 300nM of each reverse primer, and 100nM of each probe, with 4μL of DNA sample. The optimal cycling conditions were determined to be: initial denaturation of 3min @ 95°C followed by 40 cycles of 15s @ 95°C and 30s @ 61°C. All assays achieved an efficiency between 91.4% and 100.5%, except for Cy5 quantification in the first and third of 3 total 96-well plates used in the study. These assays achieved an efficiency of 124.4% and 138.8%, respectively. Efficiency was calculated using a diluted DNA standard (10-fold dilution starting with 2ng/mL) for each strain.

### ^1^H nuclear magnetic resonance spectroscopy-based metabolomics

Samples were prepared for ^1^H NMR spectroscopy as described by Dona et al. [53]. Samples were thawed at room temperature and centrifuged at 12,000 xg at 4°C for 10 minutes, before 540 μl of supernatant was combined with 60 μl of buffer (pH 7.4; 1.5mM KH2PO4, 0.1% TSP (3-(trimethylsilyl)propionic-2,2,3,3-d4 acid sodium salt) in 100% D2O) and transferred to a SampleJet NMR tube (Bruker BioSpin, Rheinstetten, Germany). Standard one-dimensional (1D) ^1^H-NMR spectra with water pre-saturation were acquired at 300 K using a 600 MHz Avance III spectrometer (Bruker), equipped with a SampleJet autosampler (Bruker). A total of 32 scans were collected into 64,000 data points for each sample. Spectra were automatically phased, baseline corrected and calibrated to the TSP resonance at δ^1^H 0 in Topspin 3.1 software (Bruker). The spectra were imported into MATLAB R2014a (The Mathworks, Inc., Natick, MA, USA). Biologically irrelevant regions of the spectra were removed (TSP resonance at δ^1^H 0 and residual water peak δ^1^H 4.5- 5.2) before peak alignment by recursive segment-wise peak alignment (RSPA) [54]. The loadings of pairwise principal component analysis models, as well as manual comparisons between fresh media spectra and spent media of each bacterial strain, were used to identify metabolites generated or consumed in each experiment. To further identify metabolites that may only have been produced or consumed in co-culture, group means for all 21 co-culture conditions were compared to fresh media across the entire spectra. The relevant regions of the spectra were integrated to calculate relative spectral intensities for each metabolite. Metabolite identities were assigned by reference to known spectra in multiple databases. For all analyses, integrals were centered by subtracting the mean value for each metabolite in the blank samples, then scaled by the maximum absolute value of all centered values (so that the minimum and maximum possible scaled values for each metabolite were -1 and +1, respectively). The peak integral values, scaled values, peak integration regions and identities, and associated R code for analysis and visualization is available in the git repository. Raw spectra are currently being submitted to the Metabolights database [55].

### Differential abundance testing

DNA quantification data for each sample group (each strain in monoculture and each unique co-culture condition) were tested for normality using the Shapiro-Wilk test (implemented via the shapiro.test function in R version 3.4.2) [56]. The data for all but one monoculture sample groups were normally distributed, but the majority of co-culture sample groups were non-normally distributed, so the non-parametric Mann-Whitney *U*-test was chosen to test for differential DNA abundance. The same procedure was performed for the metabolomic data, and the majority of sample groups and metabolites were found to be non-normally distributed, so the Mann-Whitney U-test was performed to test for differential metabolite abundance as well, identifying metabolites as either produced, consumed, or unchanged based on testing results and the value of the group mean relative to the fresh media. For tests of differential abundance, the false discovery rate (FDR) was controlled using the Benjamini-Hochberg procedure [57]. For DNA differential abundance testing, the number of sample groups used for FDR control was 21 (6 monocultures and 15 co-cultures). For metabolite differential abundance testing, the number of sample groups used for FDR control was 1806 (21 mono- and co-culture groups, each with 86 integrated metabolite peaks that were tested for differential abundance against fresh media samples). The results of all normality and differential abundance testing, and notebooks performing the calculations, are available in the github repository accompanying this work.

### Constant Yield Expectation (ConYE) model

For each sample group, metabolite integrals were centered by subtracting the mean value of the metabolite integral in fresh medium (i.e. negative control). Centered integrals were then scaled by the max of the absolute values across all sample groups for each metabolite, resulting in values for each metabolite being scaled between -1 and +1, with at least one sample group taking a value of -1 or +1 for each metabolite. For each monoculture sample group, the mean of each scaled, centered metabolite was then divided by the mean DNA abundance of the corresponding strain, resulting in a metabolite yield specifying the amount of increase or decrease of a metabolite per unit of DNA for each strain. For each co-culture sample, the expected concentration of each metabolite was determined by multiplying the abundance of each strain in co-culture by the monoculture-derived yield, summing the two quantities from each strain, then subtracting the concentration of the metabolite in the fresh medium using the mean across negative controls (N=9). Using the calculated expected concentration from all samples within a co-culture group, deviation from expectation was determined by comparing expected concentrations to the observed concentrations in co-culture. Differential abundance was determined using the Mann-Whitney l/-test with FDR control via the Benjamini-Hochberg procedure [57] and the mean of differences between expected and observed concentrations were recorded. The sample size used for FDR control was 1290 (15 co-culture groups, each compared to a simulated ConYE value for each of 86 metabolites). Notebooks performing the calculations for ConYE, and the results of all tests, are available in the github repository accompanying this work.

### Draft genome-scale metabolic network reconstruction

Draft genome-scale metabolic network reconstructions (GENREs) were generated for ASF356, ASF360, ASF361, ASF492, ASF500, ASF502 (not included in the present study), and ASF519 using a local installation of ProbModelSEED [34, 58], with draft genome sequences for the strains from the same experimental stock used in this study [59]. Briefly, ProbModelSEED annotates the genome for each organism using RAST [60], which in turn identifies metabolic functions associated with genes or sets of genes. This process results in a draft GENRE containing high-confidence reactions for each species. To enable biomass production in the GENRE, gapfilling is performed with uptake enabled for any metabolite with a transporter annotated in the draft GENRE (i.e. simulating a rich medium). The resulting GENRE contains the original reactions associated with the organism’s annotated genome, as well as non-gene associated reactions added to enable biomass production. ProbModelSEED also assigns reaction probabilities to each reaction that can be added during gapfilling, which are derived using sequence similarity for genes which did not meet the similarity threshold for annotation via RAST. These probabilities are incorporated during gapfilling, leading to preferential addition of reactions for which there was some genetic evidence.

### Metabolomics-constrained gapfilling

After generation of draft GENREs, we added functionality to the GENREs using a previously-generated supernatant metabolomics dataset in which the same ASF strains were grown in the same medium used in this study [21]. Using the original metabolite annotations for the dataset, we constrained the GENRE for each ASF strain by forcing production and consumption of any metabolite with a z-score normalized abundance of > +1 or < -1, respectively. This was enforced by setting the lower bound of the exchange reaction for produced metabolites to 0.001 mmol/(g dry weight * hour), and the upper bound for exchange reactions for consumed metabolites to -0.001 mmol/(g dry weight * hour). The value of the constraint (0.001 mmol/(g dry weight * hour)) was chosen to be arbitrarily-low, since absolute changes in metabolite concentrations were not derived in the metabolomics dataset used for gapfilling. Then, we set the remaining boundary conditions for each GENRE to represent the medium in which they were grown (as described in the *in silico* simulations section), and forced arbitrarily low flux through the biomass reaction (0.005/hour). Then, we checked for transporters for each metabolite for each strain that enabled import (for consumed metabolites) and export (for produced metabolites). If a suitable transport reaction was not present, we added a transporter from the ModelSEED biochemistry database and constrained the directionality to be as observed (e.g. import only for consumed metabolites, export only for produced metabolites). Transporter assignments are provided in Supplemental Table S2. We then performed gapfilling using a modified version of the SMILEY algorithm [61] as implemented in Cobrapy v0.5.11. Algorithmic details are provided in Supplementary text. We used the set of all ProbModelSEED reactions for which reaction probabilities were assigned as the universal reactions for gapfilling, except for reactions including O_2_. We weighed the penalty for addition of each of these reactions by 1 - *p*, where *p* is the reaction probability, which ranges from 0 for unlikely reactions to 1 for highly likely reactions. The effect of this penalty is that high-probability reactions are assigned lower penalties, and are thus more likely to be added to the GENRE during gapfilling. For each ASF strain, metabolomics-constrained gapfilling was performed 10 times, each for 10 iterations, resulting in an ensemble of 100 GENREs. All 100 GENREs for each strain were unique (i.e. none of the iterations resulted in identical reaction sets being added to the draft GENRE).

### GENRE quality control

All GENREs within the ensemble for each strain were assessed for mass balance. To perform this assessment, an intracellular demand reaction was added for each metabolite in the GENRE, and all exchange reactions were closed. Flux through each demand reaction was then optimized iteratively, and demand reactions that could carry flux indicated presence of a mass-imbalanced reaction that allowed spontaneous generation of the metabolite. This process identified three reactions in the draft GENREs that were mass-imbalanced (SEED ids: rxn15543 in ASF519 GENRE; rxn33894 and rxn30984 in ASF361). These reactions were removed from the draft GENREs, and the metabolomics-constrained gapfilling process was repeated using the draft GENREs with the reactions removed. Since the draft reconstructions were generated in October 2016, these reactions have since been removed from the ModelSEED biochemistry database.

All GENREs within the ensemble for each strain were also assessed for infeasible ATP production, an issue commonly identified in draft-quality GENREs [62]. Using boundary conditions that mimic the *in vitro* medium (as in *in silico* simulations, below), we optimized flux through an ATP demand reaction, and found that all GENREs for all strains generated between 0.5 and 1.9 mmol ATP/(g dry weight * hour). Normalizing this value by the uptake of lactose, which was 0.22 mmol/(g dry weight * hour) for all strains, gives a yield range of 2.27-8.64 units of ATP per unit of lactose, well within reason for anaerobic organisms (for example, *Escherichia coli* is known to produce 2.2-3.2 units of ATP per unit of glucose when grown anaerobically) [63]. Although erroneous energy generating cycles may be present in the GENREs presented here, the realistic ATP yield determined for all GENREs suggests that they do not influence ATP production in this particular media condition, and are thus unlikely to influence simulation results.

### *In silico* simulations

Flux balance analysis (FBA) was performed using version 0.8.1 of the cobrapy package [64]. Ensembles of GENREs were analyzed using Cobrapy methods through the Medusa package (unpublished, https://github.com/gregmedlock/Medusa/). Media composition was determined by calculating exact concentrations for defined supplements (Hemin, Vitamin K1, Lactose, Tween-20), and a concentration of 1mM was assumed to allow an uptake rate of 1 mmol/(g dry weight * hour). For media components with approximately known concentrations in LB (amino acids), the uptake rate was set to 5 mmol/(g dry weight * hour) based on a concentration of around 5 mM for most amino acids in LB [65]. For components detected via metabolomics that were not amino acids or supplemented, and therefore likely originated from the yeast extract in LB, the maximal uptake rate was set to 0.1mmol/(g dry weight * hour). For *in silico* media supplements and knockouts, a metabolite was considered essential if removal of the metabolite from the *in silico* medium caused the flux through biomass to fall below 1 E-5/hour (used because of limits of numerical precision for the solvers used; use of a lower threshold (1 E-10/hr) does not affect these results).

### Code and data availability

All raw and processed data and all code used in this project except software used to process raw NMR spectra are availability at https://github.com/gregmedlock/asf_interactions. Where possible, Jupyter notebooks [66] are used for reproducibility and to display results alongside corresponding analyses.

## Acknowledgements

We thank Michael Wannemuehler and Gregory Phillips for providing ASF strains. We acknowledge the University of Virginia Advanced Research Computing Services staff for assistance in setting up software used for gapfilling on the University of Virginia high-performance computing cluster. We thank Jie Liu for helpful guidance on hydrolysis probe design for qPCR, and Thomas Moutinho for experimental assistance performing sample extraction. We thank all members of the Papin lab for helpful project feedback.

References

## Supplementary Text

**Supplemental Table S1.** Sequences for primers, probes, and amplified products, and fluorophore and quencher pairs for each probe.

**Supplemental Table S2.** Transport reactions added to the GENRE for each species.

## Metabolomics-constrained gapfilling

Metabolomics-constrained gapfilling was performed to ensure the GENRE for each species could produce biomass in the *in vitro* medium and produce and consume metabolites as determined by supernatant metabolomics. We used a modified version of the growmatch algorithm [61] with variable reaction penalties calculated in ProbModelSEED [58]. We implemented and applied the modified version of our algorithm in cobrapy v0.5.11 [64]. The algorithm is formally defined as:

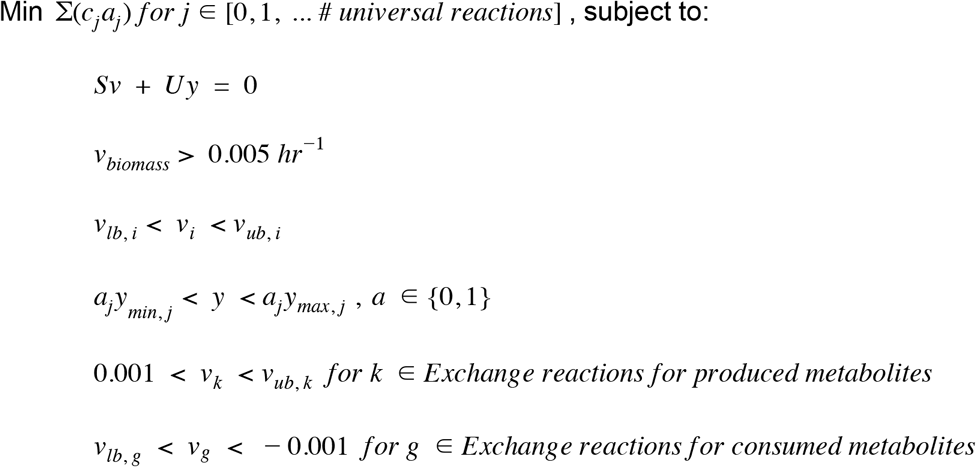

Where *c* = 1 –*p* and is the reaction cost associated with including each reaction from the universal reaction bag, *p* is the probability of each reaction (derived from sequence information using ProbModelSEED; reactions not assigned a probability receive a probability of 0) and *a* is the integer indicator for each reaction *j* in the universal reaction bag used during gapfilling (here, we use the ModelSEED biochemistry database). *S* is the stoichiometric matrix, *v* is the vector of flues through each reaction represented in the stoichiometric matrix, *U* is the universal reaction bag, *y* is the vector of fluxes through reactions in the universal reaction bag (which is multiplied by the integer *a* to force flux to take a zero or non-zero value), *v_biomass_* is flux through the biomass reaction (which we constrain to take a minimum value of 0.005 *hr*^−1^ to force an arbitrarily low, non-zero amount of growth), *v_k_* are the fluxes through exchange reactions for metabolites that were designated as produced, and *v_g_* are the fluxes through exchange reactions for metabolites that were designated as consumed. *v_lb_*, the lower bound of flux through a reaction, was 0 for irreversible reactions and -1000 for reversible reactions. *v_ub_*, the upper bound of flux through a reaction, was 1000 for all reactions. *v_lb_* for exchange reactions were set to -1000 for all metabolites detected in the medium by NMR spectroscopy and 0 for metabolites not detected. We performed gapfilling for 10 independent runs for each species, in which each run had 10 dependent iterations that each generate a solution containing a set of reactions that, when added to the GENRE and activated, satisfy the constraints (all metabolites can be produced and consumed as indicated, and biomass can be produced). Within each run, the penalty for each reaction was increased by setting *c* = 2*c* to encourage unique solutions. For reactions in the ModelSEED biochemistry that did not receive probabilities because they have no associated gene (e.g. spontaneous reactions), we set *c* = 100 to discourage addition of these reactions unless they were essential for any solution to be found. After each of 10 independent runs, reaction penalties were reset to their original values prior to beginning the next run. We reduced the integrality threshold in cobrapy to 1E-8 from the original value of 1E-6, because the default setting returned many solutions that did not meet the constraints applied due to numerical error for ASF361 (e.g. the reaction list returned did not actually enable biomass production for this species because reactions from the universal reaction bag had values for *y* that were below 1E-6; decreasing the integrality threshold properly returned these reactions). For every ASF strain, all 100 GENREs constructed were unique.

